# Geographic assessment of cancer genome profiling studies

**DOI:** 10.1101/827683

**Authors:** Paula Carrio Cordo, Elise Acheson, Qingyao Huang, Michael Baudis

**Affiliations:** Institute of Molecular Life Science, University of Zurich, Winterthurerstrasse 190, 8057, Zurich, Switzerland; SIB, Swiss Institute of Bioinformatics, Winterthurerstrasse 190, 8057, Zurich, Switzerland; Department of Geography, University of Zurich, Switzerland

**Keywords:** Cancer genomics, online resource, geodata, population assignment, meta-data, genome profiling

## Abstract

Cancers arise from the accumulation of somatic genome mutations, which can be influenced by inherited genomic variants and external factors such as environmental or lifestyle-related exposure. Due to the heterogeneity of cancers, precise information about the genomic composition of germline and malignant tissues has to be correlated with morphological, clinical and extrinsic features to advance medical knowledge and treatment options. With global differences in cancer frequencies and disease types, geographic data is of importance to understand the interplay between genetic ancestry and environmental influence in cancer incidence, progression and treatment outcome.

In this study, we analysed the current landscape of oncogenomic screening publications for geographic information content and quality, to address underrepresented study populations and thereby to fill prominent gaps in our understanding of interactions between somatic variations, population genetics and environmental factors in oncogenesis. We conclude that while the use of proxy derived geographic annotations can be useful for coarse-grained associations, the study of geo-correlated factors in cancer causation and progression will benefit from standardized geographic provenance annotations. Additionally, publication derived geographic provenance data allowed us to highlight stark inequality in the geographies of cancer genome profiling, with a near lack of sizeable studies from Africa and other large regions.

## Introduction

Cancer is one of the top causes of mortality globally, and the understanding of its genesis and pathophysiological mechanisms remains one of the major challenges in life sciences. Although the last decades have witnessed major improvements in cancer diagnostics and therapeutics - partially driven by rapid advances in genomic screening techniques such as DNA array and sequencing technologies - the overall spectrum of genomic alterations in malignant transformation and progression remains poorly understood.

As a multistep genomic disease, different factors influence the transformation of somatic cells into malignant clones. Whereas the majority of oncogenomic variants arise as mutations in the DNA of somatic cells and affect genes involved in proliferation, differentiation and control of apoptosis, inherited (“germline”) genomic variants can predispose to specific malignancies (1, 2). As external modifiers of the individual cancer risk, environmental factors such as pollution levels, intensity of UV radiation or exposure to infectious agents have been found to contribute to carcinogenetic processes to a varying extent. While environmental or lifestyle-mediated exposure related to micro- or macrogeographic origin can promote disparities in incidence and clinical outcome in individuals (3, 4), a less well-defined contribution is provided through ancestry-specific biases in the occurrence of cancer-promoting genome variants (5). Well known examples here are the population specific enrichment of BRCA1 gene variants in persons of Ashkenazi jewish ancestry compared to mixed reference populations (6); the higher somatic mutation frequencies for TP53, EP300, and NFE2L2 in Chinese patients from suffering from esophageal squamous cell carcinoma compared to “Caucasian” patients (7), or the significant molecular differences existing between prostate cancers in “African Americans” vs. “Caucasicans” (SPINK1 over-expression, ERG rearrangement and PTEN deletion are less frequent in the first group)(8).

Since highly variable cancer rates by ethnicity and geography have been observed for multiple cancer types (9), inclusion of geographic and population background information in cancer related data analysis projects should be considered for improving the understanding of environmental and population related contributions. Indeed, strong support for the systematic inclusion of geographic and population parameters in cancer data analyses comes from esophageal squamous cell carcinoma (ESCC), a cancer type with a high incidence in China compared to Europe and America, including some areas of remarkably frequent occurrence such as Chaoshan. To understand the contributions of environmental, genetic and cultural risk factors, studies in this particular disease have looked at familial correlation, genetic susceptibility (polymorphism for some chemical metabolizing genes involved with ESCC), lifestyle factors (alcohol, cigarette, tea and coffee consumption), environmental factors due to geographic variations (levels of selenium, strontium or zinc and dietary preference for consumption of fermented fish and preserved fish sauce), and relation with human papillomavirus infection (10). However, current patient-derived oncological models predominantly lack population data and tend to overrepresent “Caucasian” individuals (11). Given the current lack of standardized annotations for geographic sample provenance in prominent genome data collections such as NCBI’s Gene Expression Omnibus (GEO (12); www.ncbi.nlm.nih.gov/geo/) or EBI’s ArrayExpress (www.ebi.ac.uk/microarray-as/aer/), an exploration of published cancer genome studies using a geographic approximation method could provide a general picture of study geographies, help to understand the utility of proxy data and provide arguments about structured “geodata” annotations in biomedical research.

Cancer genomics data collections - representing the results of data curated from scientific studies and primary data resources - are essential for advancing our understanding of molecular mechanisms and guiding the improvement of treatment protocols. Whole-genome profiling for structural and sequence variations has become an integral part of the molecular assessment of cancer samples and *in-vivo* systems and has led to thousands of publications of original studies. The corpus of scientific literature related to cancer genome screening experiments, pre-selected through similarities in subject and general scope, offers an opportunity for the analysis of longitudinal directions and possible gaps in this specific field of biomedical research.

Progenetix is a publicly available cancer genome data resource (*progenetix.org*), originally established in 2001 (13) and maintained through our group at the University of Zurich UZH and Swiss Institute of Bioinformatics SIB. The original goals of the resource were to make copy number variation (CNV) data from Comparative Genomic Hybridization (CGH (14, 15)) studies comparable across different cancer types, and to counteract biases from extrapolations based on small-batch experimental results. Through the continuous compilation of oncogenomic profiling publications since 1993 (first clinical CGH samples) up to now using a standard set of query parameters against NCBI’s PubMed database (16), Progenetix’s literature collection provides a representative - though certainly not exhaustive - view of the cancer genomics publication landscape, currently containing 3’240 curated articles focusing on primary publications of individual cancer studies (excluding reviews and technical papers). By intent, it covers published data analyzed by array and chromosomal Comparative Genomic Hybridization experiments (aCGH including SNP arrays (17, 18); cCGH), as well as Whole Genome or Whole Exome Sequencing (WGS, WES (19, 20)) studies. For a subset of 1’547 articles, the database contains a total of 93’640 sample-specific genomic profiles which allow for comparative meta-analyses of genomic copy number imbalance profiles across more than 400 diagnostic classes.

Here we present an overview of the global distribution of cancer studies based on the literature data collected for the Progenetix database, used as an example for a curated knowledge resource covering the corpus of publications in a defined research area and containing readily available geographic provenance information. We highlight the importance of geographic data (populations and environmental) and aim for the improvement of standards and consensus in metadata annotation of relevant clinical and individual features which would help elucidate the multi-factorial nature of cancer through epidemiological and correlation studies.

## Materials and Methods

### Data description

As result of the standard Progenetix literature curation process 3’240 publications were collected and included in the Progenetix database. The publications, spanning the years 1993 to 2019, were retrieved from PubMed by combining keywords using boolean AND (intersection) or boolean OR (union) such as: ((“whole genome sequencing” OR WGS) OR (“whole exome sequencing” OR WES) OR (array AND genomic) OR (SNP AND array) OR “comparative genomic hybridization” OR CGH OR aCGH) AND (cancer OR leukemia OR lymphoma). For each publication, the number of analysed biosamples (i.e. instance of tumor material profiled) was determined, separately for the main genome screening technologies (WGS, WES, aCGH, cCGH). Geographic point coordinates were assigned to each set of samples using the address (city) of the first author of the associated publication, serving as a proxy for the samples’ origins. Point coordinates were obtained for each city name using the external geographic database GeoNames (www.geonames.org).

In contrast to the publication collection of the Progenetix resource, individual genome profiling experiments are labeled after review of existing information from associated publications or repository information for their “best available” geographic origin, using a precedence of *sample specific data* >*experiment location* >*first author proxy*, where available.

### Sample location evaluation

For a randomly selected subset of 200 articles, the article contents were manually examined in order to identify any text describing the geographic origin of the samples or patients. These locations, if present, could then be compared to the city of the first authors’ institutions for each article in this subset.

### Map representation

A kernel density raster was generated from the geographic point coordinates using the ArcGIS Kernel Density tool, using a search radius of 200km and a cell size of 25×25km, with each point weighted by the number of samples associated to that location. The final map shows a smoothed representation of the density of genome profiling experiments per square kilometer around the world.

### Network of collaborations

Author collaborations were visualized at the country level using a set of 3’093 publications from the Progenetix database where author affiliation countries could be automatically determined. Author affiliations were extracted by running the CERMINE tool (21) over the set of publication PDFs to obtain an XML file for each publication including structured metadata.

The set of affiliation countries (provided as country codes) was then extracted from these XML files for each article. We counted how many times a country appeared at least once in an article (*country counts*) and how many times pairs of countries appeared together on an article (*collaboration counts*). We generated a network graph representation of this data using the *Gephi* visualization software (22), with countries as nodes and collaborations between countries as edges.

## Results

### Sample location evaluation and proxy by first author location

Based on the manual examination of 200 publications for sample/patient location information in text, it was found that 30.5% of the articles contained explicit sample location information (e.g. city and country name) and 28.0% contained indirect sample location information (e.g. hospital name). The remaining 41.5% of the articles had no sample location information.

Regarding the agreement between sample locations and first author locations, for the 117 articles which contained sample location, 52.1% agreed on a city/province level, additional 22.2% on a country and 13.9% on a continent level. For the remainder of the articles (11.9%), the first author location or full author location list provided an incomplete or misleading picture of the sample/patient locations described in the text: 7.7% of the articles contained samples from more than one continent and in 4.2% of the articles the sample location was on a different continent to the first author location.

In summary, author proxy information provided a reasonable assignment at a “population” scale, with 74.3% of correct attribution to at least country level (with additional per-sample contributions from the “mixed provenance” articles). Nevertheless, when the attribution of geographic sample provenance is extrapolated from article authorship, there is a risk of potential non-alignment of the studies’ place of execution and the origin of the study material at higher resolution.

### Geographies of published studies

Fig 1 shows a map of the global geographic distribution of biosamples in the Progenetix database, using the first author affiliation location as a proxy for the sample location. As can be seen, the distribution is highly uneven and concentrated in a few regions of the world, in particular Western Europe and the North-Eastern United States. In Europe, the cities associated with the most number of samples are Amsterdam, London and Paris, whereas in the United States the top cities are Boston/Cambridge, New York City and San Francisco.

**Fig. 1.**
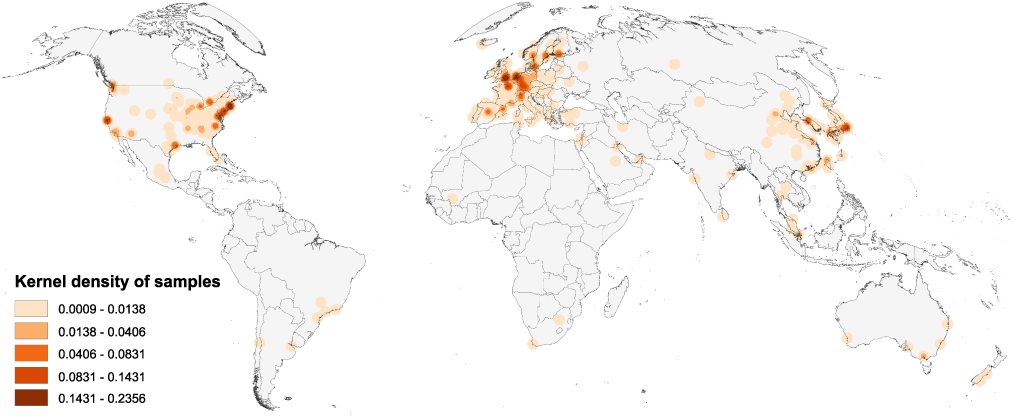
Map of the geographic distribution of genome screening experiments using the first author affiliation location proxy, derived from 3’240 publications registered in the Progenetix database, including 104’543 genomic array, 36’766 chromosomal CGH and 15’409 whole genome/exome based cancer genome profiles. The map is rendered in the Goode Homolosine Land equal area projection (kernel density per square kilometer).

Fig 2 shows the cumulative numbers of published oncogenomic screening experiments as collected in the Progenetix database. The top 10 sample-contributing countries are the United States (49’901 samples registered at Progenetix), followed by Germany (17’447), the United Kingdom (12’843), Japan (11’538), France (9’612), the Netherlands (7’777), Sweden (6’896), Spain (6’080), Canada (5’862), and China (4’937). The ranking by number of genomic arrays per year is relatively consistent over time for the top countries, with few relative shifts.

**Fig. 2.**
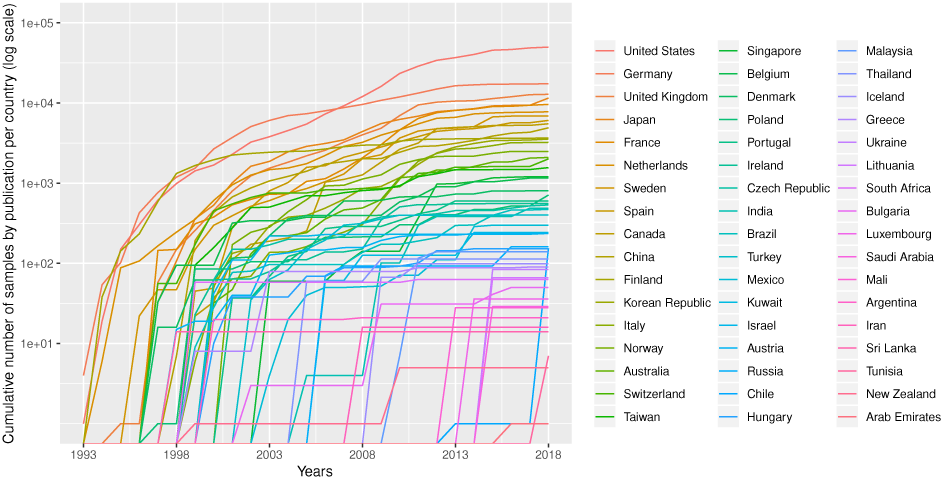
Cumulative number (in logarithmic scale) of genomic array samples (CCGH, ACGH, WES or WGS) contained in 3’240 publications registered in the Progenetix database, split by their associated country, from 1993 to 2019 (publication year).

### International collaborations

In Fig 3, we present a network graph showing the collaborations between countries based on the subset of publications (3’093) from the Progenetix database where author affiliation countries could be automatically extracted. In total, 67 countries appeared at least once in an author affiliation, with the most frequent country by far being the United States, with contributing authors on 1’215 publications (39%). The second most frequent country is Germany (523), followed by the United Kingdom (375), then the Netherlands (265) and Japan (243). 34% of the collection featured more than one country in the affiliation list, supporting a high degree of international collaboration in cancer genome research. The remaining 66% of publications featured just a single country, but with potentially many institutions collaborating. 38 publications featured more than 5 different countries in the author affiliation list, including a publication (23) with contributors from 13 different countries, the maximum in our data.

**Fig. 3.**
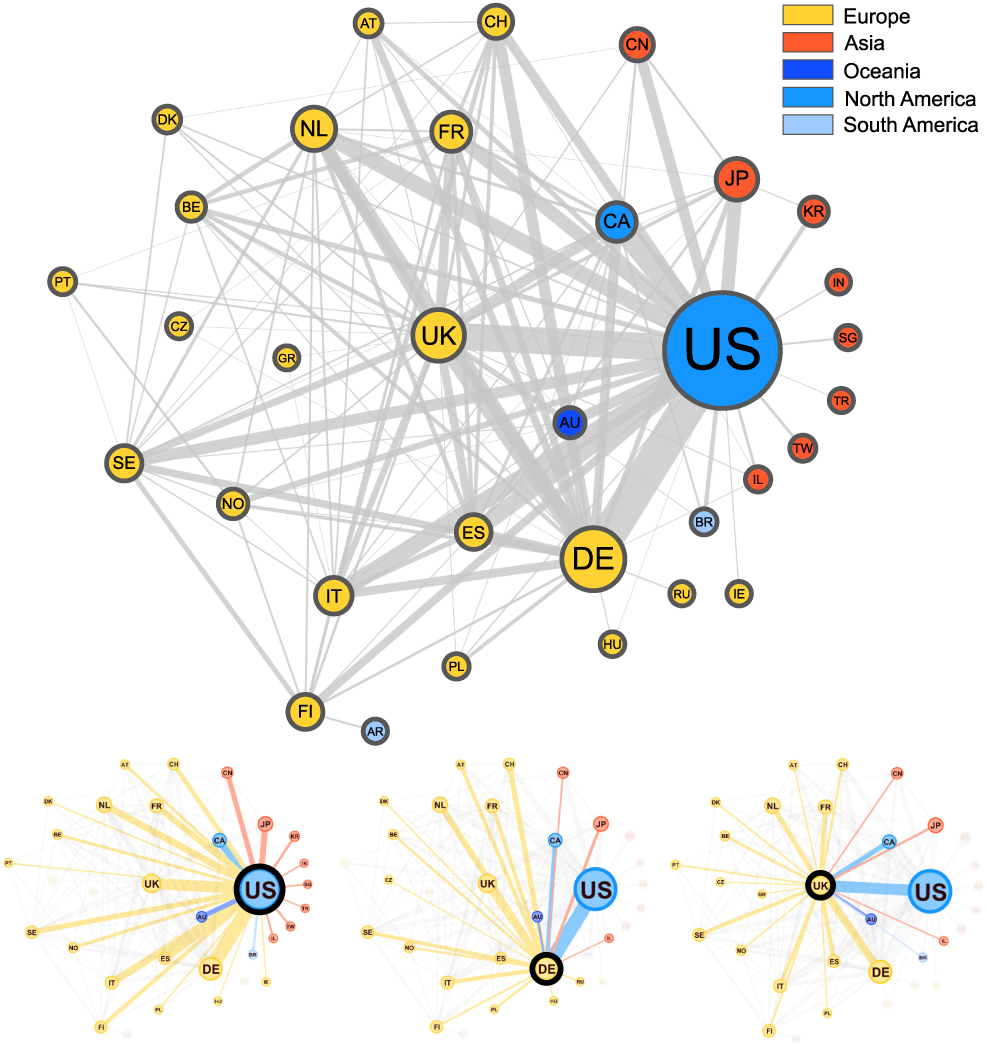
Graphical representation of the collaborations between countries based on 3’093 publications contained in the Progenetix database. Collaborating countries for the 3 most frequent countries - the United States (US), Germany (DE) and the United Kingdom (UK) - are also shown in their own graphs. Node size is proportional to the *country counts* (linearly scaled between 10 and 50), node color represents the country’s continent, and edge thickness is proportional to the *collaboration counts*. To reduce clutter in the graph, only countries appearing on more than 7 publications appear in the graph, meaning our graph depicts about 49% of all nodes and 29% of all edges in our data.

The top two most frequent country collaborations are between researchers from the US and Germany (collaborating on 122 publications), and from the US with the UK (95 collaborations), reflecting an alignment of overall country contribution frequencies and international collaborations. The US features in 6 of the next 7 most common pairs, collaborating most often with Canada (79), Italy (60), Spain (60), the Netherlands (58), Japan (49), and France (49). Though no African countries are represented in the graph, 7 of these countries appear in our data, but below the country count cut-off value (7), with Egypt appearing as an affiliation country on 5 publications, the most for the African continent.

## Standards for geographic attribution of biosamples

Geographic provenance data can be a powerful tool for genomic data stratification and exploration 4. However, as we have observed projects are frequently highly collaborative, with international contributions seen in at least one third of published studies based on the contributing authors’ affiliations. Therefore, the use of submission location as a proxy for the geolocation of the studies’ samples does not represent a rigorous procedure but rather is an operational simplification. Correlating geographic - and population background related - patient information with genomic characteristics calls for a more ambitious approach, for which standards for the geographic attribution of individuals, biosamples and technical procedures should be established.

One of the current initiatives to address challenges in the generation and - particularly - accessibility of genomic data in biomedical research and medical practice on an international scale is the Global Alliance for Genomics and Health (GA4GH) (24). A specific aim of this organisation is to allow flexible, federated data access to genomic and health-related information across national boundaries. GA4GH members pursue this goal through the development and promotion of data sharing standards and technologies, driven through working groups (e.g. for genomic knowledge standards, data security, data discovery or clinical and phenotypic data capture) and driver projects such as the ELIXIR Beacon project (25) (*beacon-project.io*).

The responsible sharing of genomics and health-related data needs to be supported by well-designed data schemas and APIs. For several of those schemas, the inclusion of geolocation parameters has been found beneficial for supporting an effective implementation of the derived protocols, e.g. for the attribution of the geographical origin of individuals and biosamples, as well as for technical provenance tracking and regulatory procedures. One model of the different domains for geolocation attribution is presented in Fig 5, where we provide a schematic representation of geolocation data objects and their use in the framework of a GA4GH schema derived, hierarchical “Individual - Biosample - Callset” model. Following this schema, the distinction between the place of residence/origin of the patient, the place where the biosample was extracted (e.g. hospital) and the facility/laboratory where the samples were sequenced or analyzed becomes feasible for different data analysis scenarios. Additionally to these geographic parameters, explicit ancestry information (e.g. using the HANCESTRO ontology; www.ebi.ac.uk/ols/ontologies/hancestro/) could provide an additional qualifier for ancestry - environment stratification. If implemented throughout an international ecosystem of GA4GH stakeholders such structured annotations will allow for diverse causative and co-relational studies with great impact on genotype/phenotype correlations and epidemiological knowledge.

**Fig. 4.**
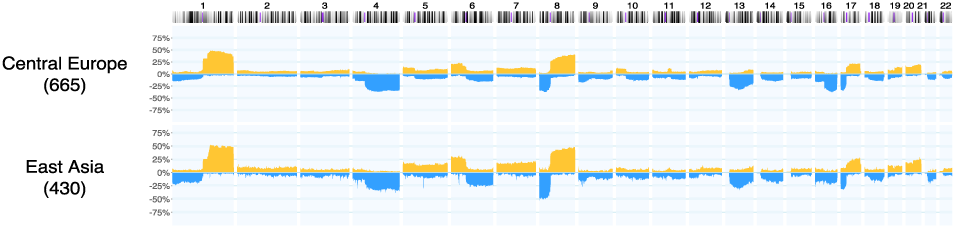
Example for a geographic subset generation, given existing geolocation data. In Hepatocellular carcinoma (ICD-O 3 Morphology 8170/3) datasets from the Progenetix collection, a query for experiments with a geographic provenance within 2’000 km from Heidelberg or Taipei, respectively, was used to generate CNV frequencies for samples from Central Europe (665 samples) and East Asia (430 samples) *ad hoc*. Such use cases are supported through the biosample-level location attributes in the (GA4GH derived) Progenetix dataschema. This example does not emphasize differences or similarities between the groups, but rather highlights the power of “one click” geographic stratification options.

**Fig. 5.**
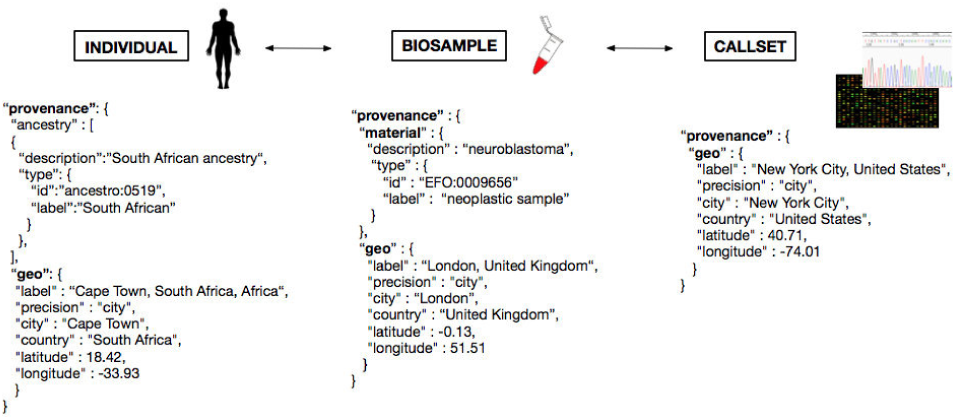
Metadata representation for a geolocation attribution approach based on concepts beingthe current schema developed at GA4GH. Each extracted “Biosample” from an “Individual”, and its “Callset” (i.e. experimental read-out) maintain a relation; ethnicity and geographical provenance of a biosample could be tracked through the attributes at the “Individual” level.

While the extension of metadata annotations with geographic provenance information will provide opportunities for better stratification of data especially in large-scale meta-analyses, possible risks of e.g. easier re-identifications of individuals through combination of genomic and location data (26) has to be considered. A careful design and review of data schemas and transmission protocols can be instrumental in lowering risks and improving the acceptance of geographic metadata annotations.

Aside from GA4GH, another important initiative within life sciences aiming at the FAIRification of data (which should follow the four foundational principles — Findability, Accessibility, Interoperability, and Reusability (27)) is bioschemas.org. It aims primarily at the annotation of data resources and repositories for easy identification of data of interest, in the biomedical domain. In the area of collaborative studies, the “ICGC ARGO” project (icgc-argo.org) is an expansion of the previous International Cancer Genome Consortium (ICGC) collaboration, which aims specifically for the inclusion of population-adjusted samples and also emphasizes the assessment of rich metadata. For the improvement of healthcare delivery systems, it intends to link past and new genomic data to clinical and health information (e.g. lifestyle parameters, patient history, diagnosis and treatment response).

## Discussion

Based on the data from 3’240 articles curated for the Progenetix resource, our analysis delivers an overview of the geographic provenance “landscape” of cancer genome profiling publications, specifically of those containing whole-genome profile from hybridization (chromosomal CGH, genomic array) and sequencing (WES, WGS) experiments. When using the city of the first author’s institution as proxy, we can show prominent disparities for geographic sample origins on a (sub-)continental scale, with only very limited contributions from studies on African, Central Asian or South American origin. While this result could be biased through the incomplete registration of relevant studies or country-specific biases in the registration of such studies in PubMed, it corresponds to the “visibility” of this research to the international research community.

Previous studies have analyzed collaborations in science using author affiliations to analyze the spatial aspects of research and their role (28, 29). Other analyses have been focused on specific regions, such as Southeast Asia (30), to analyze the collaboration networks of biomedical research and improve the understanding of local health systems. In the area of cancer research, focused efforts have lead to the creation of interactive tools such as the Global Oncology Map (*gcpm.globalonc.org*), a project which aims to promote research projects and facilitate collaborations, where users can browse and search for projects, people, and events displayed on a world map (31).

Over the last years, large collaborative initiatives have emerged with the aim of defining the genome of different cancers. With over 88 contributing projects, the ICGC has mapped the structural aberrations of 26 major cancer types across 16 countries and the European Union. Furthermore, The Cancer Genome Atlas (TCGA), with an overview of 33 different cancers based on 11’315 cases, provides a rich collection of high quality tumor samples. However, with respect to reflecting population and geographic heterogeneity, based on samples from 14 contributing countries, the representation of population diversity is nevertheless limited with a great majority of samples labeled as being from “white” individuals (8’186 samples), and other groups represented in much lower numbers - 1’325 “not reported”, 934 “black or African American”, 675 “Asian”, 27 “American Indian or Alaska native” and 13 “native Hawaiian or other Pacific Islander”. The data from the respective TCGA publications has been included in our analysis.

Numerous scientific studies have pointed out disparities in cancer incidence, prevalence and mortality based on geographic location, and called for further study of the influence of different genetic backgrounds on cancer development (8, 32, 33). A first, important step to study the influence of geography-associated (genetic, environmental, socio-economic) factors on cancer predisposition, incidence and finally biology would be to associate each analyzed cancer sample with its geographical provenance. Unfortunately, currently available repositories for genomic data lack submission protocols in which a biosample’s geographic provenance or individual’s ethnicity must be provided, and even lack requirements for a minimal consistent representation through e.g. {city, country} pairs. Therefore, most of the time the only geographic location available is based on the city where a given project has been submitted, either to a data repository or as part of the publication process. However, as seen from the manual analysis of the 200 publications, this submitter affiliation does not always reflect the geographic provenance of the biosample: A biopsy could have been taken at a hospital, then analyzed later on at a research center in a different city, country, or even continent. This leaves a great challenge in deciding whether such locations are meaningful as proxies for the genetic and/or environmental backgrounds.

Even with existing geographic provenance, such information can only provide an approximation of a patients ancestry, with varying probability and specificity depending on regional patterns of migration and admixture. For (cancer) genome profiling data with accessible genotyping information (e.g. SNP arrays, genome-wide NGS data), a recent approach of direct genetic inference allows for the assignment of population groups from shared genomic ancestry (34). While this method provides genome based access to the ancestry component of the sample provenance with superior information compared to metadata for that specific domain, it will be restricted to the subset of samples with accessible genotyping data and therefore be limited by lack of widespread data deposition and also privacy concerns associated with genotype data access. Additionally, such a genome-based approach cannot account for additional data dimensions (environmental, technical) for which geographic attribution might be beneficial.

## Conclusions

Correct attribution of the geographic origin of a patient can provide valuable information as a proxy for two general classes of factors with a known role in cancer development: A) environmental and lifestyle factors with exposure related to local or regional geographic origin, and B) ancestry related variation in occurrence or frequency of genomic variants providing heritable contributions to cancer development. However, the lack of standards and deposition requirements for geographic metadata in genomic data repositories and as part of scientific publication procedures currently limits potential benefits of geographic attribution in cancer genome studies. While the use of proxy information (e.g. location of author or data submitter) can be valuable in geographic epistemology and for coarse-grained geographic associations, detailed studies on the impact of geo-correlated factors in cancer causation and progression could be enabled through precise annotation of geographic provenance using standardized protocols and data formats.

In our study, we present a GA4GH derived concept for the structured annotation of geographic provenance data which accommodates for different levels of data attribution in genome - and possibly other - profiling analyses, and demonstrate its implementation in our “Progenetix” cancer genome profiling collection, for stratification of CNV datasets according to their large-granular geographic origin.

While the limited availability of structured geographic provenance data severely restricts options for detailed associations between geographic data and genomic parameters in cancer meta-analyses, the use of author based proxy data allows an estimation of the overall international study landscape in the field of cancer genomics. Here, our analysis of 3240 publications shows large inequalities, with the majority of studies being contributed from European, North American and East Asian groups or collaborations, and the near complete lack of accessible cancer genome profiling data from Africa and, to a lesser extent, central Asia and South America. The lack of ethnic diversity in the current data collections - reflected in these extreme global imbalances - highlights the need to properly address global disparities in cancer research to enable the study of the influence of genetic background on cancer development.

## Supplementary Information

Additional information and supplementary data can be found in the online repository (github.com/progenetix/publications: *2020-Carrio-Cordo-cancergeographies*).

## Acknowledgments

We thank current and former members of the Baudisgroup at UZH for their continuing efforts in making cancer genome data accessible, and the manuscript’s reviewers for the helpful comments.

## References

1. D Malkin, FP Li, LC Strong, JF Fraumeni, CE Nelson, DH Kim, J Kassel, MA Gryka, FZ Bischoff, and MA Tainsky. Germ line p53 mutations in a familial syndrome of breast cancer, sarcomas, and other neoplasms. Science, 250(4985):1233–1238, 1990.

2. PA Futreal, Q Liu, D Shattuck-Eidens, C Cochran, K Harshman, S Tavtigian, LM Bennett, A Haugen-Strano, J Swensen, and Y Miki. Brca1 mutations in primary breast and ovarian carcinomas. Science, 266(5182):120–122, 1994.

3. RL Siegel, KD Miller, and A Jemal. Cancer statistics, 2017. CA Cancer J Clin, 67(1):7–30, 2017.

4. G Danaei, S Vander Hoorn, AD Lopez, CJ Murray, M Ezzati, and Risk Assessment collaborating group (Cancers Comparative. Causes of cancer in the world: comparative risk assessment of nine behavioural and environmental risk factors. Lancet, 366(9499):1784–1793, 2005.

5. DS Tan, TS Mok, and TR Rebbeck. Cancer genomics: Diversity and disparity across ethnicity and geography. J Clin Oncol, 34(1):91â–101, 2016.

6. CI Szabo and MC King. Population genetics of brca1 and brca2. Am J Hum Genet, 60(5): 1013–1020, 1997.

7. Jiaying Deng, Hu Chen, Daizhan Zhou, Junhua Zhang, Yun Chen, Qi Liu, Dashan Ai, Hanting Zhu, Li Chu, Wenjia Ren, Xiaofei Zhang, Yi Xia, Menghong Sun, Huiwen Zhang, Jun Li, Xinxin Peng, Liang Li, Leng Han, Hui Lin, Xiujun Cai, Jiaqing Xiang, Shufeng Chen, Yihua Sun, Yawei Zhang, Jie Zhang, Haiquan Chen, Shijian Zhang, Yi Zhao, Yun Liu, Han Liang, and Kuaile Zhao. Comparative genomic analysis of esophageal squamous cell carcinoma between asian and caucasian patient populations. Nature Communications, 8(1): 1533, 2017. ISSN 2041-1723. doi: 10.1038/s41467-017-01730-x.

8. Francesca Khani, Juan Miguel Mosquera, Kyung Park, Mirjam Blattner, Catherine O’Reilly, Theresa Y MacDonald, Zhengming Chen, Abhishek Srivastava, Ashutosh K Tewari, Christopher E Barbieri, et al. Evidence for molecular differences in prostate cancer between african american and caucasian men. Clinical Cancer Research, 2014.

9. Wensheng Zhang, Andrea Edwards, Erik K. Flemington, and Kun Zhang. Racial disparities in patient survival and tumor mutation burden, and the association between tumor mutation burden and cancer incidence rate. Scientific Reports, 7(1):13639, 2017. ISSN 2045-2322. doi: 10.1038/s41598-017-13091-y.

10. WR Tang, ZJ Chen, Kun Lin, Min Su, and WW Au. Development of esophageal cancer in chaoshan region, china: association with environmental, genetic and cultural factors. International journal of hygiene and environmental health, 218(1):12–18, 2015.

11. Santiago Guerrero, Andrés López-Cortés, Alberto Indacochea, Jennyfer M García-Cárdenas, Ana Karina Zambrano, Alejandro Cabrera-Andrade, Patricia Guevara-Ramírez, Diana Abigail González, Paola E Leone, and César Paz-y Miño. Analysis of racial/ethnic representation in select basic and applied cancer research studies. Scientific reports, 8, 2018.

12. R Edgar, M Domrachev, and AE Lash. Gene expression omnibus: Ncbi gene expression and hybridization array data repository. Nucleic Acids Res, 30(1):207–210, 2002.

13. M Baudis and ML Cleary. Progenetix.net: an online repository for molecular cytogenetic aberration data. Bioinformatics, 17(12):1228–1229, 2001.

14. A Kallioniemi, OP Kallioniemi, D Sudar, D Rutovitz, JW Gray, F Waldman, and D Pinkel. Comparative genomic hybridization for molecular cytogenetic analysis of solid tumors. Science, 5083(258):818–821, 1992.

15. S Joos, H Scherthan, MR Speicher, J Schlegel, T Cremer, and P Lichter. Detection of amplified dna sequences by reverse chromosome painting using genomic tumor dna as probe. Hum Genet, 90(6):584–589, 1993.

16. H Cai, N Kumar, N Ai, S Gupta, P Rath, and M Baudis. Progenetix: 12 years of oncogenomic data curation. Nucleic Acids Res, 42(Database issue):D1055–62, 2014.

17. S Solinas-Toldo, S Lampel, S Stilgenbauer, J Nickolenko, A Benner, H Dohner, T Cremer, and P Lichter. Matrix-based comparative genomic hybridization: biochips to screen for genomic imbalances. Genes Chromosomes Cancer, 4(20):399–407, 1997.

18. X Zhao, C Li, JG Paez, K Chin, P. Jänne, TH Chen, L Girard, J Minna, D Christiani, C Leo, JW Gray, WR Sellers, and M Meyerson. An integrated view of copy number and allelic alterations in the cancer genome using single nucleotide polymorphism arrays. Cancer Res, 64(9):3060–3071, 2004.

19. TJ Ley, ER Mardis, L Ding, B Fulton, MD McLellan, K Chen, D Dooling, BH Dunford-Shore, S McGrath, M Hickenbotham, L Cook, R Abbott, DE Larson, DC Koboldt, C Pohl, S Smith, A Hawkins, S Abbott, D Locke, LW Hillier, T Miner, L Fulton, V Magrini, T Wylie, J Glasscock, J Conyers, N Sander, X Shi, JR Osborne, P Minx, D Gordon, A Chinwalla, Y Zhao, RE Ries, JE Payton, P Westervelt, MH Tomasson, M Watson, J Baty, J Ivanovich, S Heath, WD Shannon, R Nagarajan, MJ Walter, DC Link, TA Graubert, JF DiPersio, and RK Wilson. Dna sequencing of a cytogenetically normal acute myeloid leukaemia genome. Nature, 456 (7218):66–72, 2008.

20. PJ Campbell, PJ Stephens, ED Pleasance, S O’Meara, H Li, T Santarius, LA Stebbings, C Leroy, S Edkins, C Hardy, JW Teague, A Menzies, I Goodhead, DJ Turner, CM Clee, MA Quail, A Cox, C Brown, R Durbin, ME Hurles, PA Edwards, GR Bignell, MR Stratton, and PA Futreal. Identification of somatically acquired rearrangements in cancer using genome-wide massively parallel paired-end sequencing. Nat Genet, 40(6):722–729, 2008.

21. Dominika Tkaczyk, Pawel Szostek, Mateusz Fedoryszak, Piotr Jan Dendek, and Lukasz Bolikowski. CERMINE: automatic extraction of structured metadata from scientific literature. International Journal on Document Analysis and Recognition (IJDAR), 18(4):317–335, December 2015. ISSN 1433-2825. doi: 10.1007/s10032-015-0249-8.

22. Mathieu Bastian, Sebastien Heymann, and Mathieu Jacomy. Gephi: an open source software for exploring and manipulating networks. Icwsm, 8(2009):361–362, 2009.

23. I P M Tomlinson, M Dunlop, H Campbell, B Zanke, S Gallinger, T Hudson, T Koessler, P D Pharoah, I Niittymäkix, S Tuupanenx, L A Aaltonen, K Hemminki, A Lindblom, A Försti, O Sieber, L Lipton, T van Wezel, H Morreau, J T Wijnen, P Devilee, K Matsuda, Y Nakamura, S CastellvÞ-Bel, C Ruiz-Ponte, A Castells, A Carracedo, J W C Ho, P Sham, R M W Hofstra, P Vodicka, H Brenner, J Hampe, C Schafmayer, J Tepel, S Schreiber, H Völzke, M M Lerch, C A Schmidt, S Buch, V Moreno, C M Villanueva, P Peterlongo, P Radice, M M Echeverry, A Velez, L Carvajal-Carmona, R Scott, S Penegar, P Broderick, A Tenesa, and R S Houlston. Cogent (colorectal cancer genetics): an international consortium to study the role of polymorphic variation on the risk of colorectal cancer. British Journal Of Cancer, 102:447 EP –, 2009.

24. Global Alliance for Genomics and Health. Genomics. a federated ecosystem for sharing genomic, clinical data. Science, 352(6291):1278–1280, 2016.

25. M Fiume, M Cupak, S Keenan, J Rambla, S de la Torre, SOM Dyke, AJ Brookes, K Carey, D Lloyd, P Goodhand, M Haeussler, M Baudis, H Stockinger, L Dolman, I Lappalainen, J Törnroos, M Linden, JD Spalding, S Ur-Rehman, A Page, P Flicek, S Sherry, D Haussler, S Varma, G Saunders, and S Scollen. Federated discovery and sharing of genomic data using beacons. Nat Biotechnol, 37(3):220–224, 2019. doi: 10.1038/s41587-019-0046-x.

26. M Gymrek, AL McGuire, D Golan, E Halperin, and Y Erlich. Identifying personal genomes by surname inference. Science, 339(6117):321–324, 2013.

27. Mark D Wilkinson, Michel Dumontier, IJsbrand Jan Aalbersberg, Gabrielle Appleton, Myles Axton, Arie Baak, Niklas Blomberg, Jan-Willem Boiten, Luiz Bonino da Silva Santos, Philip E Bourne, et al. The fair guiding principles for scientific data management and stewardship. Scientific data, 3, 2016.

28. Koen Frenken, Sjoerd Hardeman, and Jarno Hoekman. Spatial scientometrics: Towards a cumulative research program. Journal of Informetrics, 3(3):222–232, 2009.

29. Raj Kumar Pan, Kimmo Kaski, and Santo Fortunato. World citation and collaboration networks: uncovering the role of geography in science. Scientific reports, 2:902, 2012.

30. Vincent Schubert R Malbas. Mapping the collaboration networks of biomedical research in southeast asia. Technical report, PeerJ PrePrints, 2015.

31. Edward L Trimble, Ali A Chisti, Jane A Craycroft, Kalina Duncan, Manaswi Gupta, Daniel Gutierrez, Ilyana Rosenberg, Nour Sharara, Sudha Sivaram, Hillary M Topazian, et al. Launching an interactive cancer projects map: A collaborative approach to global cancer research and program development. Journal of global oncology, 1(1):7–10, 2015.

32. Tanya Keenan, Beverly Moy, Edmund A. Mroz, Kenneth Ross, Andrzej Niemierko, James W. Rocco, Steven Isakoff, Leif W. Ellisen, and Aditya Bardia. Comparison of the genomic landscape between primary breast cancer in african american versus white women and the association of racial differences with tumor recurrence. Journal of Clinical Oncology, 33 (31):3621–3627, 2015. doi: 10.1200/JCO.2015.62.2126.

33. Daniel SW Tan, Tony SK Mok, and Timothy R Rebbeck. Cancer genomics: diversity and disparity across ethnicity and geography. Journal of Clinical Oncology, 34(1):91–101, 2015.

34. Qingyao Huang and Michael Baudis. Enabling population assignment from cancer genomes with snp2pop. bioRxiv, page 368647, 2019.

